# Genome-wide Association Study for Noise-induced Cochlear Synaptopathy

**DOI:** 10.1101/311407

**Keywords:** GWAS, HMDP, NIHL, eQTL

## Abstract

In order to elucidate the genetic architecture of the auditory hair cell synapse and the susceptibility to noise-induced cochlear synaptopathy, we are providing the first genome-wide association study with 111 strains (n=695) of the Hybrid Mouse Diversity Panel based upon the strain variation of the wave 1 P1-N1 amplitude of the auditory brainstem responses both before and after noise exposure. Based on this association analysis and our cochlear gene expression data, we identified several novel loci and prioritized positional candidate genes related to cochlear synaptopathy, especially after exposure to noise.

**Abstract:** This is the first genome-wide association study (GWAS) with the Hybrid Mouse Diversity Panel (HMDP) to define the genetic landscape of the auditory hair cell synapse and the susceptibility to noise-induced cochlear synaptopathy. We tested 5-week old female mice (n=695) from 111 HMDP strains (n= 6-7/strain) at baseline and post noise exposure using ABR wave 1 suprathreshold amplitudes (P1-N1 at 80 dB SPL) at 8, 12, 16, 24 and 32 kHz tone burst stimuli. Mice were exposed for 2 hours to 10 kHz octave band noise (OBN) at 108 dB SPL. A broad range of suprathreshold ABR wave 1 amplitude were detected across the HMDP strains. At the genome-wide significance threshold (-logP = 5.39), associations on Chr. 3 and Chr. 16 were identified at baseline. Also, association peaks on Chr. 2 and Chr. 13 were determined post noise exposure. In order to prioritize candidate genes, we generated gene expression microarray profiles using RNA isolated from cochleae in 64 HMDP strains (n =3 arrays per strain). We then used EMMA to perform an association analysis between all SNPs and array probes mapping within each region. A total of 17 genes (2 within Chr. 3 association, 6 within Chr. 2 association and 9 within Chr. 13 association) of these 3 loci were identified with at least 1 probe whose expression was regulated by a significant *cis* eQTL in the cochlea. Also, the genetic architecture of noise induced cochlear synaptopathy is distinct from that of baseline auditory nerve/synapse integrity. In summary, from this GWAS and our eQTL data, we identified 4 novel loci and prioritized positional candidate genes related to cochlear synaptopathy at baseline and after exposure to noise.

## Introduction

The sensation of hearing is the result of mechanical impulses (sound waves) being transmitted through the fluid filled cochlea and coded into neural impulses that travel along the 8^th^ cranial nerve through the brainstem to higher association centers within the cerebral cortex. There is increasing evidence that the ability to hear and understand as we age, in a variety of environments, is dictated by the neural connectivity of the hair cells within the cochlea and their contacts with auditory nerve fibers (ANFs). Recent data in both mice and humans suggests there exists a “hidden hearing loss” resulting from synaptopathy between inner hair cells (IHCs) and type-I ANFs (1). Disruption of these synapses not only leads to denervation but also a slow degeneration of spiral ganglion neurons (2).

While there are few practical ways of identifying these types of losses in the clinical setting, there is evidence in mice that cochlear synaptopathy can be captured by the amplitude of the first wave (wave I) of the auditory-evoked potential (3). In their experiments, noise exposure led to a convincing permanent decline in the suprathreshold ABR wave I amplitude despite recovery of otoacoustic emissions and auditory brainstem responses (ABR) thresholds. It has been determined that the wave I amplitude correlates strongly with the integrity of auditory synapses and spiral ganglion fibers (4).

Recently, evidence for this “hidden hearing loss” has been demonstrated in human temporal bone specimens. (5). It appears that noise exposure over time results both pre- and postsynaptic ribbon loss. The loss of synapses between hair cells and their respective auditory nerve can lead to great difficulty hearing in noise (6).

Age-related hearing loss (ARHL) has been shown to result from loss of sensory cells and neurons (7). It has been shown in mice that ARHL results from a similar synaptic loss between IHCs and SGNs and the cochlear synaptopathy also precedes hair cell loss and threshold shift in the aging mouse ear (8).

Our laboratory has recently described age-related and noise-induced threshold shifts in 100 strains of the Hybrid Mouse Diversity Panel (HMDP) (9) and, with these data, we have begun to define the genetic landscape of this complex trait (10) (11). In an effort to begin to explore the genetic landscape of synaptopathy, we are reporting the first genome-wide association study (GWAS) in mice, corrected for population structure, based upon the strain variation of the wave 1 amplitude of the ABR both at baseline and after noise exposure.

## Materials and Methods

### Ethics statement

The Institutional Care and Use Committee (IACUC) at University of Southern California endorsed the animal protocol for the Hybrid Mouse Diversity Panel (HMDP) inbred strains (IACUC 12033). Strains and genotypes are accessible from Jackson Labs (www.jax.org). All results required to corroborate the conclusions presented here are provided entirely within this article.

### Hybrid Mouse Diversity Panel Strains and Genotypes

5-week old female mice (n=695) from 111 Hybrid Mouse Diversity Panel strains (n= 6-7/strain) were purchased from Jackson Laboratories. A detailed characterization of the HMDP is provided in Bennett BJ, et al. 2010 (12). Mice were aged until 5 weeks and accommodated in sterilized cages with autoclaved food and water.

The phenotypes in the HMDP strains were managed using genotypes of 500,000 single nucleotide polymorphisms (SNPs) obtained from the Mouse Diversity Array (minor allele frequencies > 5%; missing genotype frequencies <10%) emerged a final collection of 200,000 SNPs.

### ABR Wave 1 P1-N1 values for wave 1 suprathreshold amplitude

In order to analyze baseline and post noise exposure auditory brainstem response wave 1 P1-N1 measurements, stainless steel electrodes were placed subcutaneously at the vertex of the head and the right mastoid, with a ground electrode at the base of the tail. Stainless steel electrodes were placed subcutaneously at the vertex of the head and the left mastoid. A ground electrode was placed at the base of the tail. Mice were anesthetized with an intraperitoneal injection of ketamine (80 mg/kg body wt) and xylazine (16 mg/kg body wt). Mouse body temperature was maintained through the use of a TCAT-2DF temperature controller and the HP-4 M heating plate (Physitemp Instruments Inc., Clifton, NJ). Artificial tear ointment was applied to the eyes and each mouse was recovered on a heating pad.

Test sounds were presented using a (Intelligent Hearing Systems) speaker attached to an 8-inch long tube that was inserted into the ear canal. Auditory signals were presented as tone pips with a rise and a fall time of 0.5 msec and a total duration of 5 msec at the frequencies 8, 12, 16, 24, and 32 kHz. Tone pips were delivered below threshold and then increased in 5 dB increments until goal of 100 dB. Signals were presented at a rate of 30/second. Responses were filtered with a 0.3-3 kHz pass-band (x10,000 times). For each stimulus intensity 512 waveforms were averaged. Data was stored for offline analysis of ABR peak-to-peak (P1-N1) values for wave 1 amplitudes at 80 dB SPL. Post-exposure thresholds were evaluated by the same method 2 weeks post-exposure.

### Noise Exposure Protocol

Using a method adapted from Kujawa and Liberman (2009) (13), 6 week old mice were exposed for 2 hours to 10 kHz octave band noise (OBN) at 108 dB SPL. The noise exposure protocol was previously described by White CH et al. (14). The cage was arranged in a soundproof chamber (MAC-1) created by Industrial Acoustics (IAC, Bronx, NY) and the sound chamber was lined with soundproofing acoustical foam. Noise recordings were performed with a speaker (Fostex FT17H Tweeter) constructed into the top of the sound chamber. Calibration of the deleterious noise was done with a B&K sound level meter with a variation (1.5 dB) over the cage.

For 2 hours, mice were positioned in a circular exposure cage with 4 shaped compartments and were capable to move about within the compartment. Testing involved the right ear only.

All hearing tests were performed in a separate (MAC-1 soundproof) chamber in order to eliminate both environmental and electrical noise. An acquisition board (National Instruments Corporation, Austin, Texas) was regulated by custom software used to generate the stimuli and to measure the responses. Stimuli were provided by a custom acoustic system (two miniature speakers, and sound pressure was measured by a condenser microphone).

### Data Analysis

We performed the association testing by FaST-LMM linear mixed model (15), a method that is capable to account for population structure.

This model was developed using the SNPs from all other chromosomes to improve power when testing all SNPs on a specific chromosome. This procedure includes the SNP being tested for association in the regression equation only once.

Genome-wide significance threshold in the HMDP was determined by the family-wise error rate (FWER) as the probability of observing one or more false positives across all SNPs per phenotype. We performed 100 different sets of permutation tests and parametric bootstrapping of size 1000 and observed that the genome-wide significance threshold at a FWER of 0.05 corresponded to P = 4.1 x 10^−6^, similar to that used in previous studies with the HMDP (12). This is approximately an order of magnitude larger than the threshold obtained by Bonferroni correction (4.6 x 10^−7^), which would be an overly conservative estimate of significance because nearby SNPs among inbred mouse strains are highly correlated with each other.

### GWAS Candidate Genes

RefSeq genes were downloaded from the UCSC genome browser (http://genome.ucsc.edu/cgi_bin/hgGateway) using the NCBI Build37 genome assembly to characterize genes located in each association.

The 95% confidence interval for the distribution of distances between the most significant and the true causal SNPs, for simulated associations that explain 5% of the variance in the HMDP, is 2.6 Mb (12). Only SNPs mapping to each associated region were used in this analysis. We selected SNPs that were variants in at least one of the HMDP CI strains.

### Gene Expression Data

Cochleae from each 8-wk-old mouse were isolated from the 64 HMDP strains (2–4 mice/strain). The inner ear was microdissected and the surrounding soft tissue and the vestibular labyrinth was removed. The dissected cochleae were then frozen in liquid nitrogen and ground to powder. RNA was extracted and purified by placing cochlea samples in RNA lysis buffer (Ambion). The sample was incubated overnight at 4°C, centrifuged (12,000 x g for 5 min) to pellet insoluble materials, and RNA isolated following manufacturer’s recommendations. This procedure generates 300 ng of total RNA per mouse.

Gene expression analysis was performed, and gene expression measurements were taken using Illumina’s mouse whole genome expression kit, BeadChips. Amplifications and hybridizations were performed according to Illumina’s protocol (Southern California Genome Consortium microarray core laboratory at University of California, Los Angeles). RNA (100 ng) was reverse transcribed to cDNA using Ambion cDNA synthesis kit (AMIL1791) and then converted to cRNA and labeled with biotin. Further, 800 ng of biotinylated cRNA product was hybridized to prepare whole genome arrays and was incubated overnight (16–20 hr) at 55°C. Arrays were washed and then stained with Cy3-labelled streptavadin. Excess stain was removed by washing and then arrays were dried and scanned on an Illumina BeadScan confocal laser scanner.

### Reagent and Data Availability

The authors state that all data necessary for confirming the conclusions presented in the article are represented fully within the article.

## Results

### Baseline and post noise exposure phenotypic variation within the HMDP

We tested 5-week old female mice (n=695) from 111 HMDP strains (n= 6-7/strain) at baseline and post noise exposure using ABR wave 1 suprathreshold amplitudes (P1- N1) at 8, 12, 16, 24 and 32 kHz tone burst stimuli.

The complete ABR wave 1 suprathreshold amplitude dataset is provided as supplemental file 1 (baseline, S1) and 2 (post noise exposure, S2).

A broad range of suprathreshold ABR wave 1 amplitudes were detected across the HMDP with differences between the lowest and the highest strains at specific ABR stimulus frequencies demonstrated in Figures 1a and 1b.

**Figure 1.**
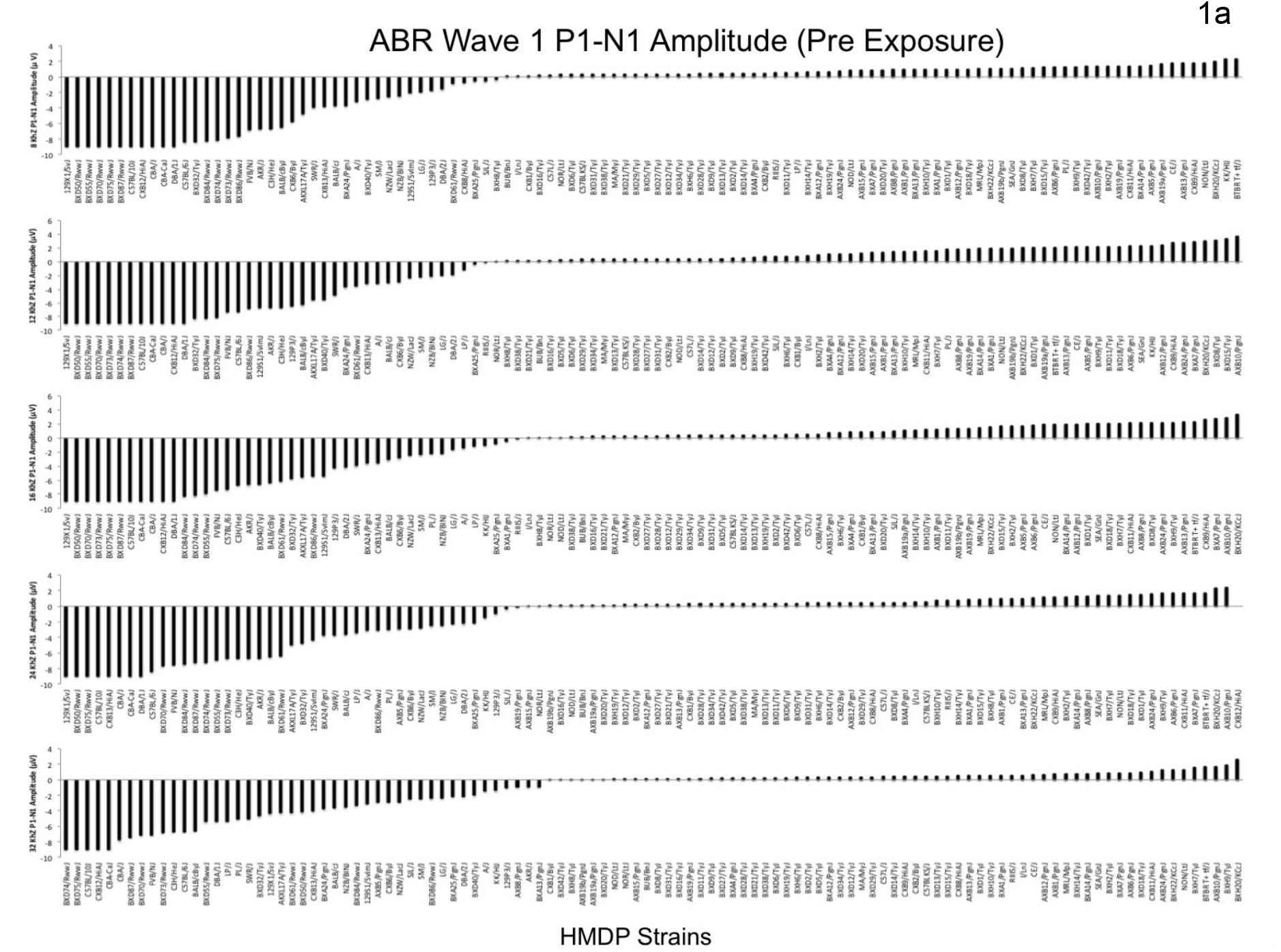
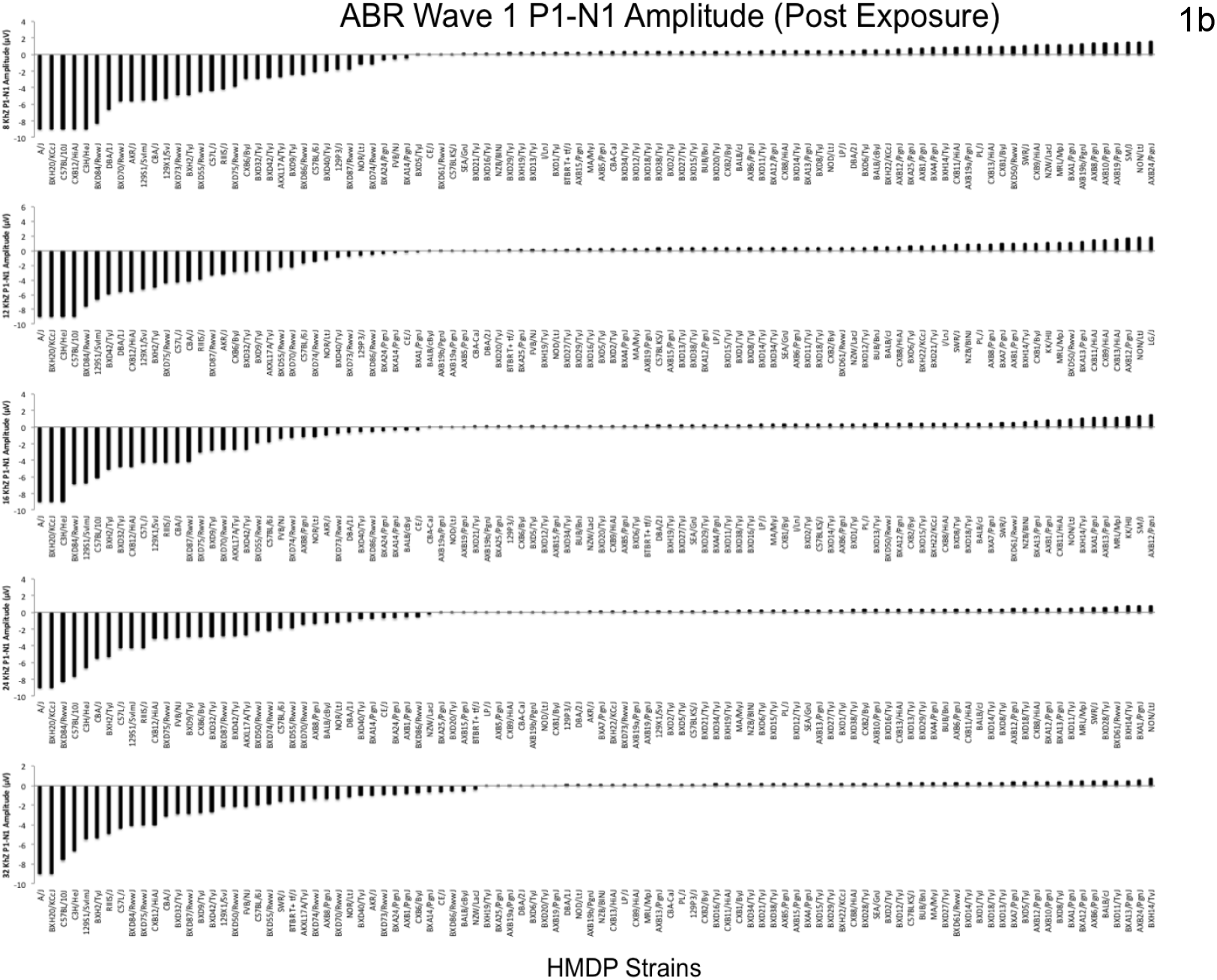
Strain Variation in ABR wave 1 amplitude. Characterization of baseline (1a) and post noise exposure (1b) P1-N1 Wave 1 amplitudes (at 80 dB SPL) in 111 HMDP inbred mouse strains.

Frequencies of 8, 12, 16, 24 and 32 kHz demonstrated differences of 3.82, 2.42, 2.62, 3.75 and 3.43-fold at baseline (Figure 1a), respectively. Frequencies of 8, 12, 16, 24 and 32 kHz demonstrated differences of 3.75, 2.43, 3.23, 5.20 and 7.5-fold post noise exposure (Figure 1b), respectively. These data supported the notion that suprathreshold ABR wave 1 amplitude variation had a genetic basis.

### GWAS for ABR P1-N1 variation at each tested frequency

Association analysis was applied to each phenotype separately to identify genetic associations for the five tone-burst stimuli. We performed a genome-wide association study for both baseline (Figure 2a and 2b) and post exposure (PTS) (Figure 2c and 2d) in order to identify loci associated at the various frequencies tested.

**Figure 2.**
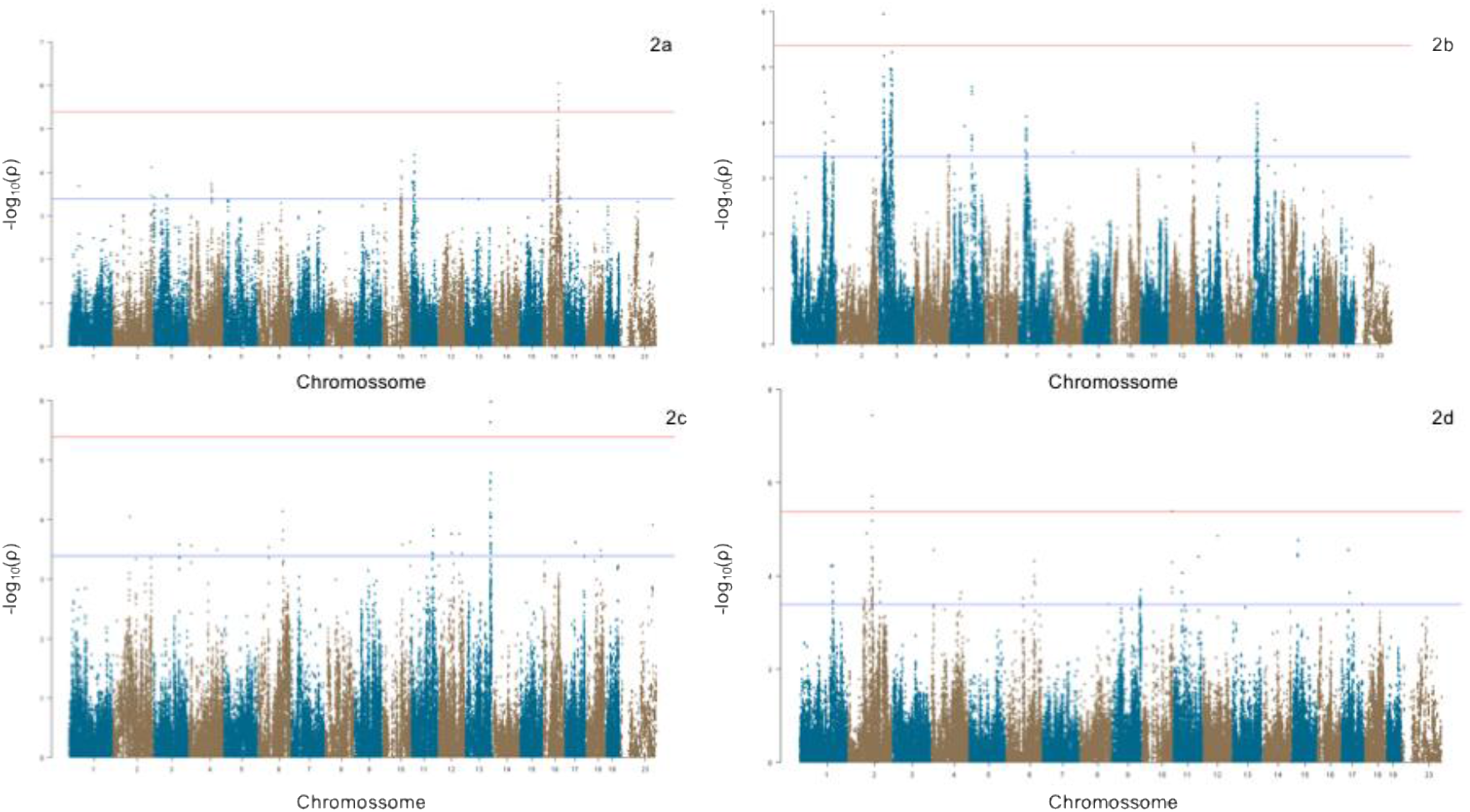
GWAS results for ABR Wave 1 P1-N1 Amplitude in the HMDP. Manhattan plot showing the association (-log10) p-values (-logP) for 16 kHz (Figure 2a) and 24 kHz (Figure 2b) at baseline and 8 kHz (Figure 2c) and 32 kHz (Figure 2d) post noise exposure in 111 HMDP inbred mouse strains. The analysis was performed using ~200,000 SNPs with a minor allele frequency > 5%. Each chromosome is plotted on the x-axis in alternating brown and blue colors. SNPs on Chr. 16, Chr. 3, Chr. 13 and Chr. 2 exceeded the predetermined genome-wide significance threshold (-logP = 5.39).

Association p-values (adjusted) were calculated for 200,000 SNPs with minor allele frequency of > 5% (p < 0.05 genome-wide equivalent for GWA using FaST-LMM in the HMDP is p=4.1 x 10^−6^, −log10P=5.39) as described above.

At the genome-wide significance threshold (-logP = 5.39), associations on Chr. 3 and Chr. 16 were identified at baseline (Figure 2). Also, association peaks on Chr. 2 and Chr. 13 were determined post noise exposure. The details of each association are provided in Table 1.

**Table 1.**
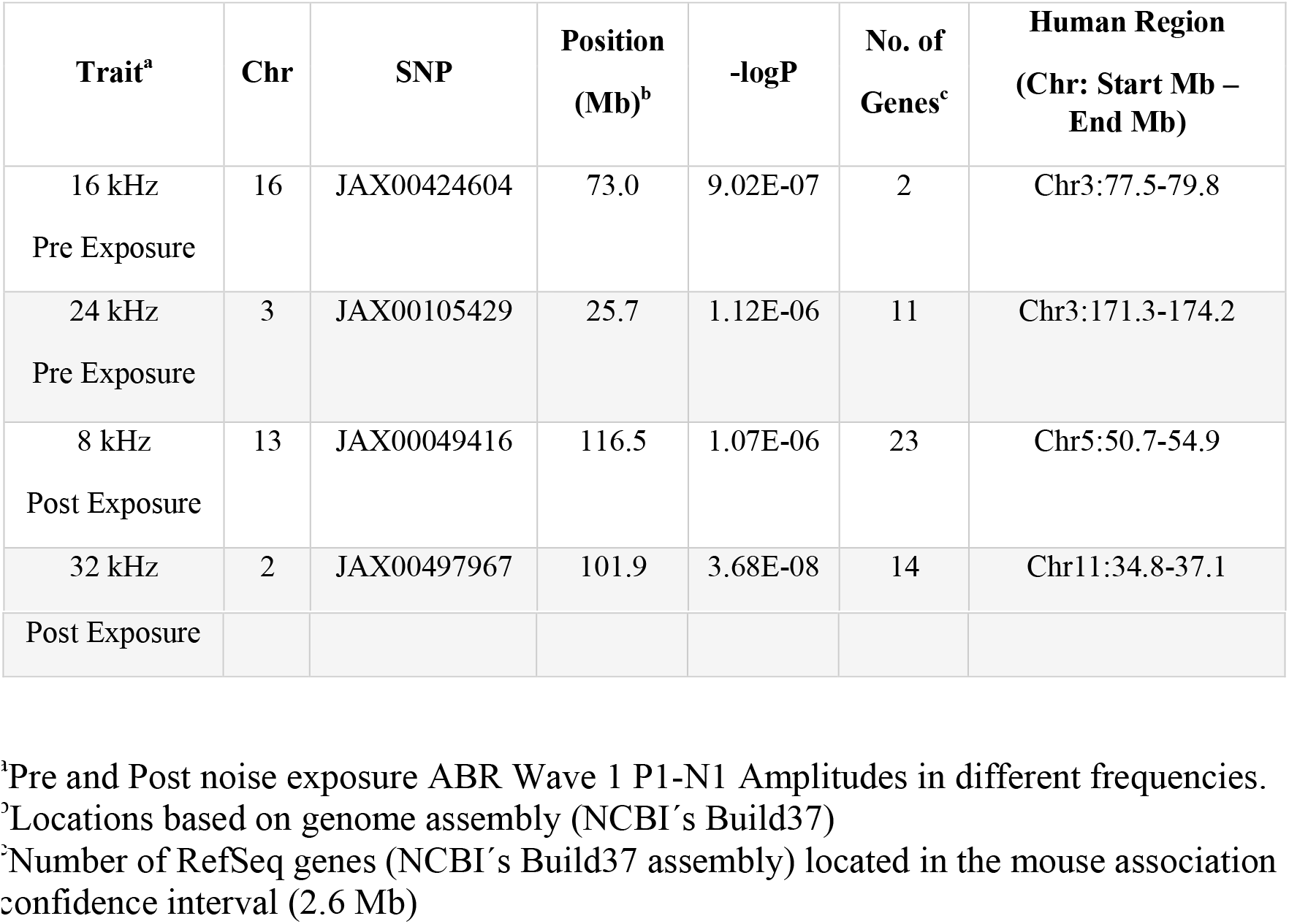
Genome-wide association results at baseline and post noise exposure.

### Association Analysis

In order to characterize these 4 associations, we defined 2.6 Mb as our interval (Table 1). At baseline, ABR wave 1 P1-N1 associations exceeding the genome-wide significance threshold were identified on chromosome 3 (Figure 3a) and 16 (Figure 3b). For post noise exposure, associations were significant on chromosome 2 (Figure 3c) and 13 (Figure 3d).

**Figure 3.**
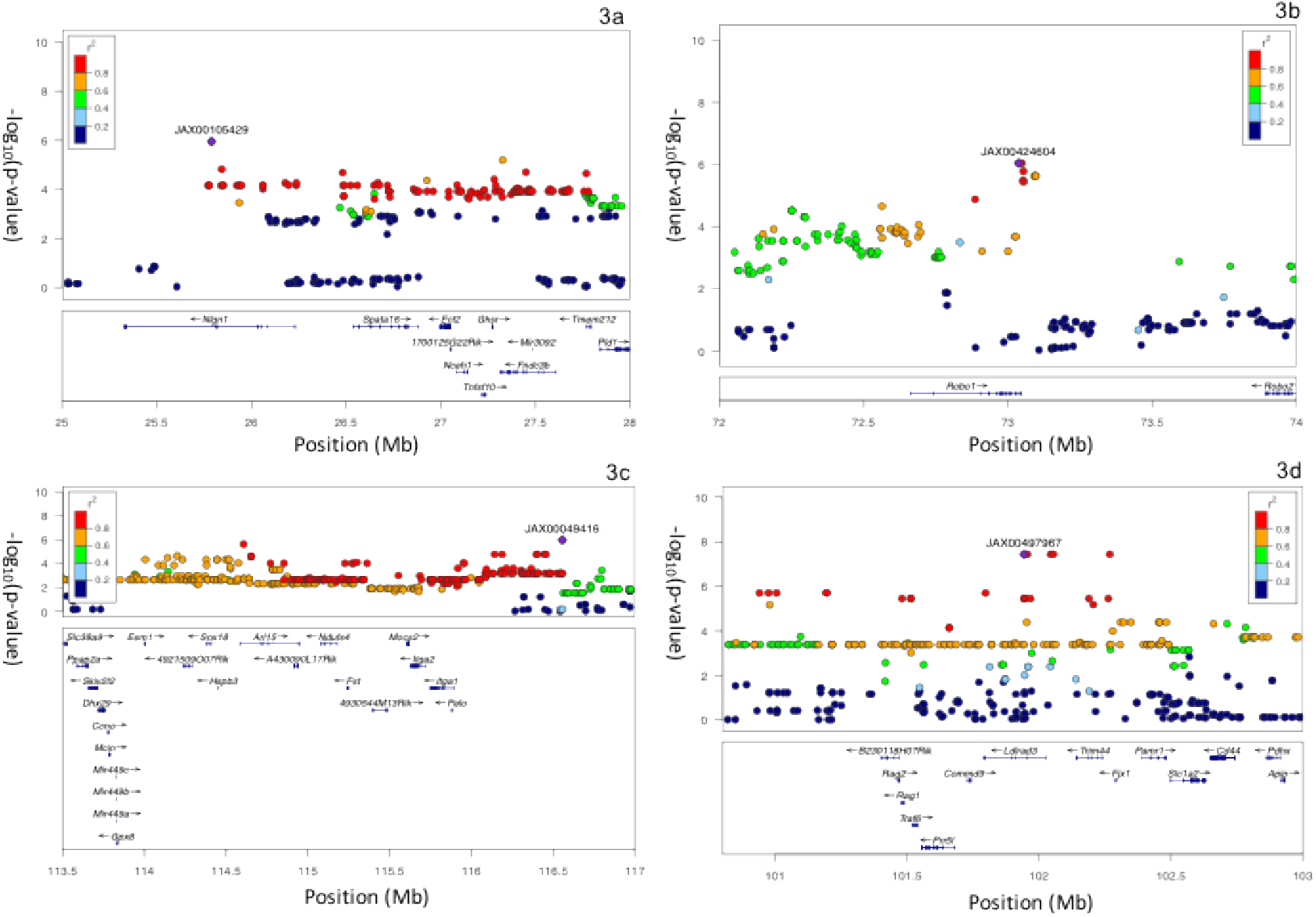
Interval of the significant association. centered on the lead SNP for 16 kHz (Figure 3a) and 24 kHz (Figure 3b) at baseline and 8 kHz (Figure 3c) and 32 kHz (Figure 3d) post noise exposure in 111 HMDP inbred mouse strains. The blue diamond represents the most significant SNP and SNPs are colored based on their LD with the most significant SNP being: red SNPs in LD at r^2^>0.8, orange SNPs in LD at r^2^>0.6 and green SNPs in LD at r^2^>0.4. The positions of all RefSeq genes are plotted using genome locations (NCBI’s Build37 genome assembly).

### Exploring Candidate Genes

Our cochlear expression data allowed us to analyze all 50 candidate genes at each association interval. Except the chromosome 16 locus, we identified genes within each of the intervals regulated by a local expression QTL (eQTL). In order to perform eQTL analysis, we generated gene expression microarray profiles using RNA isolated from cochleae in 64 HMDP strains (n =3 arrays per strain). We then used EMMA to perform an association analysis between all SNPs and array probes mapping within each region. A total of 18,138 genes were represented by at least one probe, after excluding probes that overlapped SNPs present among the classical inbred strains used in the HMDP as describe in the methods.

Loci in which peak SNPs mapped to within 2 Mb of the gene whose expression was regulated were considered “local” or cis-acting eQTLs, while SNPs mapping elsewhere were considered “distal” and presumably *trans-acting* eQTLs. We calculated the significant P-value cutoff (P = 1 x 10^−6^) for local and distal associations. These genes were prioritized based upon whether they were regulated by a local expression QTL (eQTL).

A total of 17 genes (2 within Chr. 3 association, 6 within Chr. 2 association and 9 within Chr. 13 association) of these 3 loci were identified with at least 1 probe whose expression was regulated by a significant *cis* eQTL in the cochlea (Table 2).

**Table 2.**
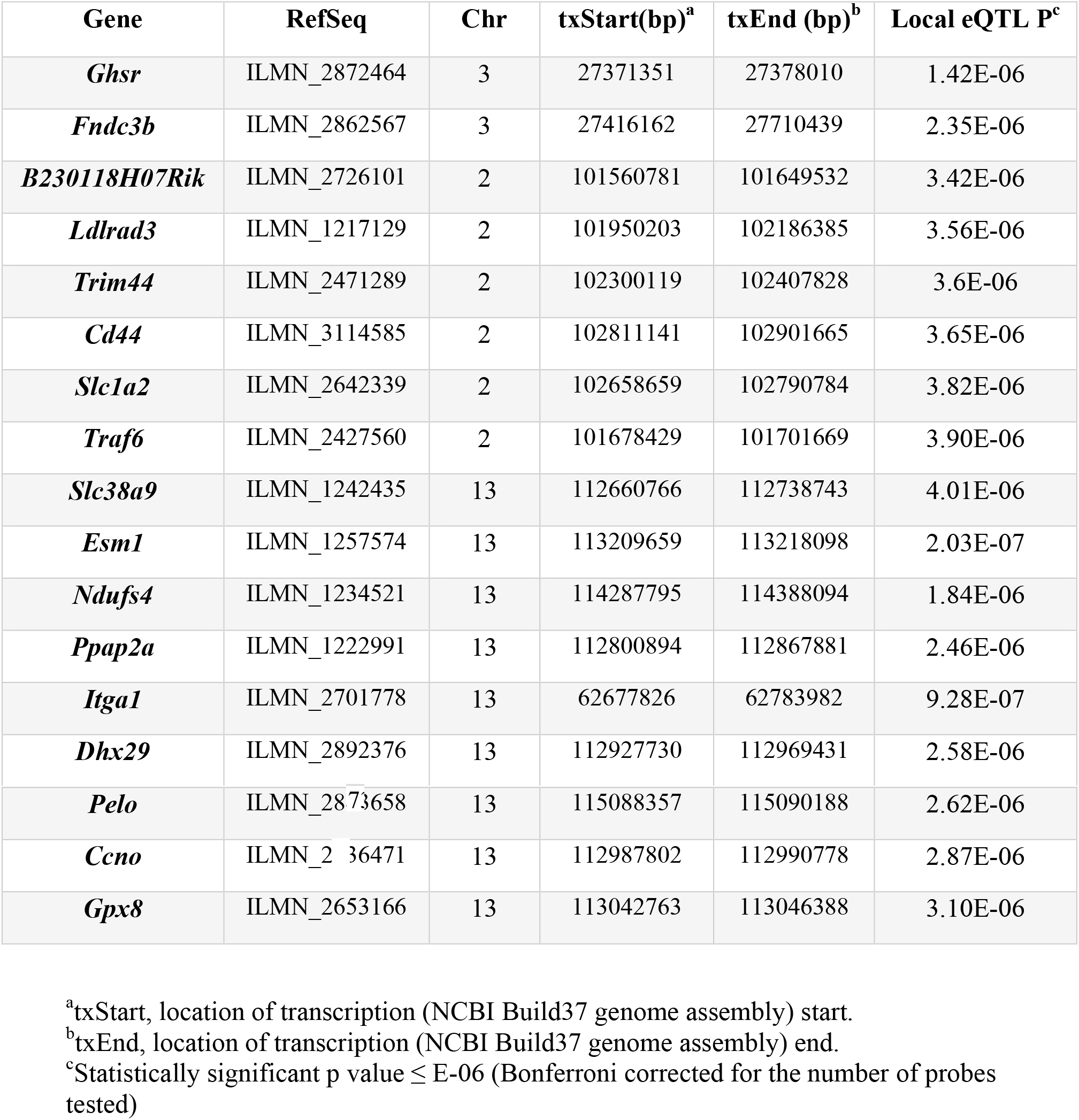
Candidate genes within 3 association peaks regulated by *cis* eQTL in the cochlea.

| Gene | RefSeq | Chr | txStart(bp)<sup>a</sup> | txEnd (bp)<sup>b</sup> | Local eQTL P<sup>c</sup> |
| --- | --- | --- | --- | --- | --- |
| *Ghsr* | ILMN_2872464 | 3 | 27371351 | 27378010 | 1.42E-06 |
| *Fndc3b* | ILMN_2862567 | 3 | 27416162 | 27710439 | 2.35E-06 |
| *B230118H07Rik* | ILMN_2726101 | 2 | 101560781 | 101649532 | 3.42E-06 |
| *Ldlrad3* | ILMN_1217129 | 2 | 101950203 | 102186385 | 3.56E-06 |
| *Trim44* | ILMN_2471289 | 2 | 102300119 | 102407828 | 3.6E-06 |
| *Cd44* | ILMN_3114585 | 2 | 102811141 | 102901665 | 3.65E-06 |
| *Slc1a2* | ILMN_2642339 | 2 | 102658659 | 102790784 | 3.82E-06 |
| *Traf6* | ILMN_2427560 | 2 | 101678429 | 101701669 | 3.90E-06 |
| *Slc38a9* | ILMN_1242435 | 13 | 112660766 | 112738743 | 4.01E-06 |
| *Esm1* | ILMN_1257574 | 13 | 113209659 | 113218098 | 2.03E-07 |
| *Ndufs4* | ILMN_1234521 | 13 | 114287795 | 114388094 | 1.84E-06 |
| *Ppap2a* | ILMN_1222991 | 13 | 112800894 | 112867881 | 2.46E-06 |
| *Itga1* | ILMN_2701778 | 13 | 62677826 | 62783982 | 9.28E-07 |
| *Dhx29* | ILMN_2892376 | 13 | 112927730 | 112969431 | 2.58E-06 |
| ***Pelo*** | ILMN_2853658 | 13 | 115088357 | 115090188 | 2.62E-06 |
| ***Ccno*** | ILMN_2736471 | 13 | 112987802 | 112990778 | 2.87E-06 |
| ***Gpx8*** | ILMN_2653166 | 13 | 113042763 | 113046388 | 3.10E-06 |

<sup>a</sup>txStart, location of transcription (NCBI Build37 genome assembly) start.
<sup>b</sup>txEnd, location of transcription (NCBI Build37 genome assembly) end.
<sup>c</sup>Statistically significant p value ≤ E-06 (Bonferroni corrected for the number of probes tested)

We then analyzed the *trans*-eQTL “hotspots.” These hotspots regulate the levels of transcripts and may disturb pathways. Also, could mediate complex gene-by-gene and gene-by-environment interactions (16).

The whole genome was divided (2-Mb bins) and the number of significant *trans* eQTL was counted in each bin. The eQTL hotspots were determined by the enrichment of gene expression traits that mapped to the same loci.

Based on this analysis, we have 5 genes regulated by the top 300 trans-eQTL “hotspots” on chromosome 2 (*B230118H07Rik, Cd44* and *Apip*) and chromosome 13 (*Esm1* and *Ndufs4*). Except *Cd44*, all these genes have no known function in the auditory system.

## Discussion

This study is the first genome-wide association study in mice using ABR wave 1 amplitude and the first large-scale assessment of this phenotype. These data will assist researchers in the study of genes and pathways involved in cochlear innervation both at baseline and after noise exposure. The HMDP has been used successfully to examine the genetics of a wide array of phenotypes by us and others including plasma lipids (17), bone density (18), blood cell traits (19), conditioned fear responses (20), gene-by-diet interactions in obesity (21), inflammatory responses (22), age-related hearing loss (10)(23), NIHL (24), diabetes (25) and heart failure (26). In many of these studies including ours, genes at the identified loci were validated as causal using engineered mouse models and several of them corresponded to loci identified in human GWAS.

Recent expansion of the depth of genotyping and the number of HMDP strains has led to an increase in the number of significant associations detected (26). There exist additional resources for the study of complex traits in mice such as the Collaborative Cross (CC), outbred rodent populations (OS), and Chromosome Substitution Strains (CSS) that have the added advantage of wild strain variation (27). There are several disadvantages that we have also considered. The CC has demonstrated breeding difficulties that would be time prohibitive. The disadvantages of the OS include the lack of available genotypes and the reproducibility advantages of inbred strains and the population structure that complicates the analysis and reduces power. The CSS, although providing substantial coverage, requires the costly generation of many rounds of subcongenic strains. The HMDP has the advantage of genomic homozygosity and complete reproducibility of phenotypic measurements and recent additions to the HMDP including the addition of ~ 75 additional strains and higher density SNP genotyping have substantially increased the power and genomic coverage (26). Although the HMDP may not capture SNP variation from wild strains, the limitations of the other resources, including the need for whole genome genotyping of successive generations in the OS and CC, the poor breeding reported for the CC, and the need for generation of subcongenics in the CSS compel us to continue with the HMDP resource.

Our GWAS generated significant associations in two regions for baseline hearing and another two separate regions post exposure to noise generating a total 50 candidate genes. Using our cochlear gene expression data and eQTL analysis we were able to narrow down candidate genes substantially. Our findings demonstrate that the genetic architecture of noise induced cochlear synaptopathy is distinct from that of baseline auditory nerve/synapse integrity.

One locus at baseline and both loci at post exposure to noise had at least two genes within the association peak regulated by a significant local cochlear eQTL. Of the 50 candidate genes at the locus, we identified 17 as having significant *cis* eQTL in the cochlea (Table 2). From these 17 candidate genes, only *Cd44* has a known function in the auditory system. *Cd44* was previously identified as preferentially expressed in the auditory sensory epithelium. Immunohistochemistry revealed that within the early postnatal organ of Corti, the expression of *Cd44* is restricted to outer pillar cells (28).

*Slc1a2* is also a strong candidate for the 32-kHz (post noise exposure) locus based on our cochlear eQTL data. *Slc1a2* encodes a member of a family of solute transporter proteins. This membrane-bound protein is the key transporter that clears the excitatory neurotransmitter glutamate from the extracellular space at synapses in the nervous system. Glutamate clearance is required for precise synaptic activation and to prevent neuronal damage from excessive activation of glutamate receptors. The synapse is composed of a presynaptic ribbon enclosed by a halo of neurotransmitter-containing vesicles within the inner hair cells (29) and a postsynaptic active zone on the cochlear nerve terminal with glutamate (AMPA-type) receptors for the released neurotransmitter (30). Although there is no known function in the auditory system, but abnormal regulation of this gene is thought to be related to several neurological disorders (31). Puel et al. (32) has shown that local application of glutamate receptor (GluR) agonists can produce dose-dependent swelling of cochlear nerve terminals contacting IHCs. The dendritic inflammation is observed under inner hair cells, but not outer hair cells, and is prevented by prior intracochlear perfusion of glutamate antagonists (33). Based on conjectures (13), this excitotoxicity would be a primary initial circumstance in the inflammatory cascade observed after noise. Thus, *Slc1a2* could play an important role in the susceptibility to noise-induced cochlear synaptopathy by regulating the glutamate clearance after exposure.

*Fst* (Follistatin), our candidate gene at the 8-kHz (post noise exposure) locus, is an antagonist of TGF-β/BMP signaling. *Fst* is expressed in the lesser epithelial ridge and at gradually higher levels close to the apex. The ascending expression of *Fst* occurs at the time of cochlear specification and is preserved throughout embryonic and postnatal development suggesting a role in cochlear tonotopy. (34).

*Neuroglin1*, a candidate gene at the 24-kHz (at baseline) locus, is a synaptic cell adhesion molecule that connects pre and postsynaptic neurons. *Neuroglin1* is responsible for signaling across the synapse which regulates synaptic activity and determines the properties of neuronal networks (35) but has no known function in the auditory system.

Our baseline GWAS data on chromosome 16 with the peak SNP within *Robo1* validates our approach. Wang SZ et al. have previously demonstrated that both *Robo1* and *Robo2* were actively expressed by spiral ganglion neurons (36). Also, in Robo1/2 double mutants at E18, spiral ganglion neurons were dislocated in the space dorsal to the cochlear epithelium and didn’t innervate hair cells. Thus, *Robo* signaling mediates spatial positioning of spiral ganglion neurons during development of cochlear. Interestingly, there were no significant alterations of the overall cochlear structure in Robo1/2 double mutants in comparison with their heterozygous littermates (hair cells persisted unaffected) (36).

Future studies will be dedicated to the validation of candidate genes through the analyses of the strains with the most extreme phenotypes and the use of transgenic and/or CRISPR models.

## Conclusions

We have performed the first comprehensive analysis of ABR wave 1 amplitude in mice and have begun to elucidate the genetic architecture of the auditory hair cell synapse and the susceptibility to noise-induced cochlear synaptopathy. We identified multiple novel loci and, using our cochlear eQTLs we prioritized positional candidate genes. These findings validate again the power of the HMDP for detecting genes related to auditory function.

